# *Aedes albopictus* (Skuse) susceptibility status to agrochemical-insecticides used in durian plantating systems in southern Thailand

**DOI:** 10.1101/2020.07.09.194746

**Authors:** Sakda Ratisupakorn, Sokchan Lorn, Nsa Dada, Aran Ngampongsai, Pawit Chaivisit, Wanapa Ritthison, Krajana Tainchum

## Abstract

High rates of dengue, chikungunya, and zika morbidity occur in southern Thailand. The intensive application of insecticides in orchards could impact not only agricultural insect pests, but also non-target insects, such as mosquitoes, or non-target beneficial insects. In this study, the population density and insecticide susceptibility of *Aedes albopictus* populations to field application concentrations of four agrochemical insecticides – cypermethrin, chlorpyrifos, carbaryl, and imidacloprid were examined. Mosquito eggs were collected from durian cultivation sites in five provinces in southern Thailand and hatched and allowed to develop to the adult stage. The study sites were categorized into three groups based on insecticide application; intensive-application of insecticides (IA), less-application of insecticides (LA), and no application of insecticides (NA). Twenty ovitraps were deployed for at least three consecutive days at each study site to collect mosquito eggs and to determine the *Ae. albopictus* population density then WHO tube assays being used to determine the susceptibility of adult mosquitoes to selected insecticides. This study represents the first report of the agrochemical insecticide susceptibility status of *Ae. albopictus* collected from durian orchards in southern Thailand. The study found that the populations of *Ae. albopictus* were susceptible to chlorpyrifos, but showed reduced mortality following exposure to lambda-cyhalothrin, carbaryl, and imidacloprid which is suggestive of the existence of resistance. These findings provide new insights into mosquito insecticide resistance focusing on *Ae. albopictus* populations and has important implications for mosquito and mosquito-borne disease control in Thailand as well as providing baseline data on which future studies can develop.

## INTRODUCTION

The viruses responsible for dengue, chikungunya, and zika are spread by mosquitoes and result in high morbidity and mortality rates every year (Thavara et al. 2009; DDC 2018). *Aedes albopictus* (Skuse) (Diptera: Culicidae), the Asian tiger mosquito, which is the vector for all these insect-borne diseases, is most commonly found in tropical and subtropical regions in both suburban and rural areas where there are open spaces with considerable vegetation (Ponlawat et al. 2005). Female *Ae. albopictus* are closely associated with activities in people’s daily life since they are present in houses and around the cultivation areas close to them, and they are common in rubber plantations as well as other tropical fruit orchards (Sullivan et al. 1971; Thammapalo et al. 2009; Tangena et al. 2016).

Tropical fruit orchards are widely cultivated in the southern region of Thailand (Tantrakonnsab and Tantrakoonsab 2018). Durian orchards are one of the most common types in southern Thailand, and numerous commercial durian growers enhance their harvest by the intensive application of agrochemical insecticides. The use of insecticides in durian orchards is especially common during off-season planting, and allows the fruits to grow gradually throughout the year. Different groups of insecticides, and different concentrations to be applied, are recommended for durian cultivation. They include; organophosphates (chlorpyrifos, methidathion), pyrethroids (lambda-cyhalothrin, cypermethrin), carbamates (carbaryl, carbosulfan), and amitraz (Wanwimolruk et al. 2015). The continuous and widespread use of agrochemical–insecticides in the durian planting system can lead to insect populations in the area, including non-target insect pests like mosquitoes (Overgaard 2006; Overgaard et al. 2005), becoming less susceptible to insecticides.

The resistance of mosquitoes to several chemicals approved for public health use has long been reported in Thailand (Chareonviriyahpap et al. 1999; Overgaard 2006; Chareonviriyaphap et al. 2013; Corbel et al. 2016). *Aedes albopictus* larvae in Phatthalung showed resistance to permethrin, while, adults in Songkhla were found to remain susceptible to deltamethrin, permethrin, fenitrothion, and propoxur (Pethuan et al. 2007) and Chuaycharoensuk et al. (2011) reported the susceptibility of adult *Ae. albopictus* from rubber plantation areas in Sadao, Songkhla to deltamethrin. Agricultural areas represent good habitats for mosquito development, and the intensive use of insecticides for crop protection and the use of other agrochemicals in those areas may contribute to the selection of insecticide resistance genes. However, mosquito populations in agricultural areas generally remain susceptible to pyrethroids, and pyrethroid-resistance does not presently pose a direct threat to vector control. Nevertheless, increased use of pyrethroids in agriculture may cause problems for vector control in future (Overgaard et al. 2005).

Because of the reported spread of insecticide resistance across different geographic locations in Thailand, an evaluation of insecticide use is needed. Moreover new insecticides which can be used as alternatives to those currently employed, and perhaps a change in the application regimens of currently used insecticides may be required to combat the threat posed by resistant mosquito strains, along with a system for monitoring the effectiveness of insecticides by local communities. Rotation systems for switching from one insecticide to another can also be designed so that the development of insecticide resistance in mosquito populations can be prevented. Cross-resistance or resistance to different insecticides approved for public health and agricultural use should also be considered when decisions are made relating to vector control.

The increasing number of dengue cases in Thailand may be in part due to failed dengue control efforts which can result from many factors other than insecticide resistance. However, in areas where insecticide resistance is a problem, the use of physiological or biological controls should be considered as an alternative to the use of insecticides (Jirakanjanakit et al. 2007a; Jirakanjanakit et al. 2007b; Pethuan et al. 2007).

Since 2016, the number of dengue cases has continued to increase, reaching high levels that have never before been recorded in southern Thailand (DDC 2018) and several hypotheses have been advanced to explain this phenomenon. These include the ineffectiveness of dengue vector control, poor self-protection against mosquito bites by those living in dengue-endemic areas, and the reduced susceptibility of mosquitoes to insecticides (Limkittikul et al. 2014). Thus, the insecticide susceptibility of *Ae. Albopictus*, which commonly breeds in orchard areas, needs to be evaluated. Some groups of insecticides, which share similar modes of action, are commonly used both by the public health authorities for vector control, as well as in durian plantations to control insect pests. Thus, the development of resistance populations to pesticides used in durian plantations in *Ae. albopictus* may lead to cross-resistance to public health insecticides. The study reported in this paper was conducted in order to investigate whether this was the case in southern Thailand. The specific objectives of the present study were to determine the density of *Ae. albopictus* in the durian planting system in southern Thailand, characterize the type and quantity of insecticides used, determine the insecticide resistance status of *Ae. albopictus* to frequently used agrochemical-insecticides in the area, and further, to characterize peoples’ attitudes to the impacts of mosquitoes and mosquito control efforts in the region.

## MATERIALS AND METHODS

### Study area

A total of 22 durian orchards in southern Thailand were surveyed and were classified by the frequency of insecticide application, as follows: intensive-application of insecticides (IA) for sites where insecticides were applied every 7-15 days (n = 12), less-application of insecticides (LA) for sites where insecticides were applied for 15 consecutive days once or twice a year (n = 3), and (NA) for sites with no application of insecticides (n = 7). The 22 durian orchards included were variously located in Chumphon (CHU), Nakhorn Si Thammarat (NAK), Phatthalung (PHA), Satun (SAT), and Songkhla (SON) provinces, and convenience sampling was used to recruit eligible participants, who were the cultivators at the orchards. Each cultivator gave permission for the study site to be accessed and samples of the mosquito eggs and immature stages to be collected. A questionnaire-based survey was then used to collect information regarding the type, frequency, and quantity of insecticides used in each orchard surveyed. Each study site was georeferenced by GPS based on its coordinates and its location was mapped using Google Maps (Figure 1). The coordinates for each location are presented in Table 1.

**Table 1.** The coordinates of the 22 durian orchards classified based on the frequency of insecticide application.

| Durian planting system | No. | Site | GPS Coordinates |
| --- | --- | --- | --- |
| **Intensive-application of insecticides (IA)** | 1 | CHU 1 | 9°52'49.9"N 98°55'04.2"E |
|  | 2 | CHU 2 | 9°52'52.7"N 98°55'07.0"E |
|  | 3 | CHU 3 | 9°53'11.8"N 98°54'35.3"E |
|  | 4 | CHU 4 | 9°53'35.0"N 98°54'47.1"E |
|  | 5 | CHU 5 | 9°53'38.4"N 98°54'50.8"E |
|  | 6 | PHA 3 | 7°39'59.8"N 99°49'57.4"E |
|  | 7 | SAT 4 | 6°52'28.8"N 99°59'58.9"E |
|  | 8 | NAK 1 | 8°48'31.6"N 99°37'36.4"E |
|  | 9 | NAK 2 | 8°48'37.7"N 99°37'27.2"N |
|  | 10 | NAK 3 | 8°48'33.5"N 99°37'28.5"E |
|  | 11 | NAK 5 | 8°44'11.1"N 99°44'15.5"E |
|  | 12 | NAK 6 | 8°46'23.2"N 99°42'58.2"E |
| **Less-application of insecticides (LA)** | 1 | NAK 4 | 8°43'59.6"N 99°44'28.4"E |
|  | 2 | SON 1 | 6°58'14.4"N 100°19'00.3"E |
|  | 3 | SON 3 | 7°00'58.0"N 100°15'28.4"E |
| **No application of insecticides (NA)** | 1 | PHA 1 | 7°40'43.9"N 99°50'18.0"E |
|  | 2 | PHA 2 | 7°40'45.0"N 99°49'56.5"E |
|  | 3 | SAT 1 | 6°54'59.2"N 99°51'19.7"E |
|  | 4 | SAT 2 | 6°54'47.0"N 99°51'24.6"E |
|  | 5 | SAT 3 | 6°47'25.8"N 100°04'46.7"E |
|  | 6 | SAT 5 | 6°52'01.2"N 100°00'23.4"E |
|  | 7 | SON 2 | 6°57'17.4"N 100°16'00.9"E |

**Figure 1.**
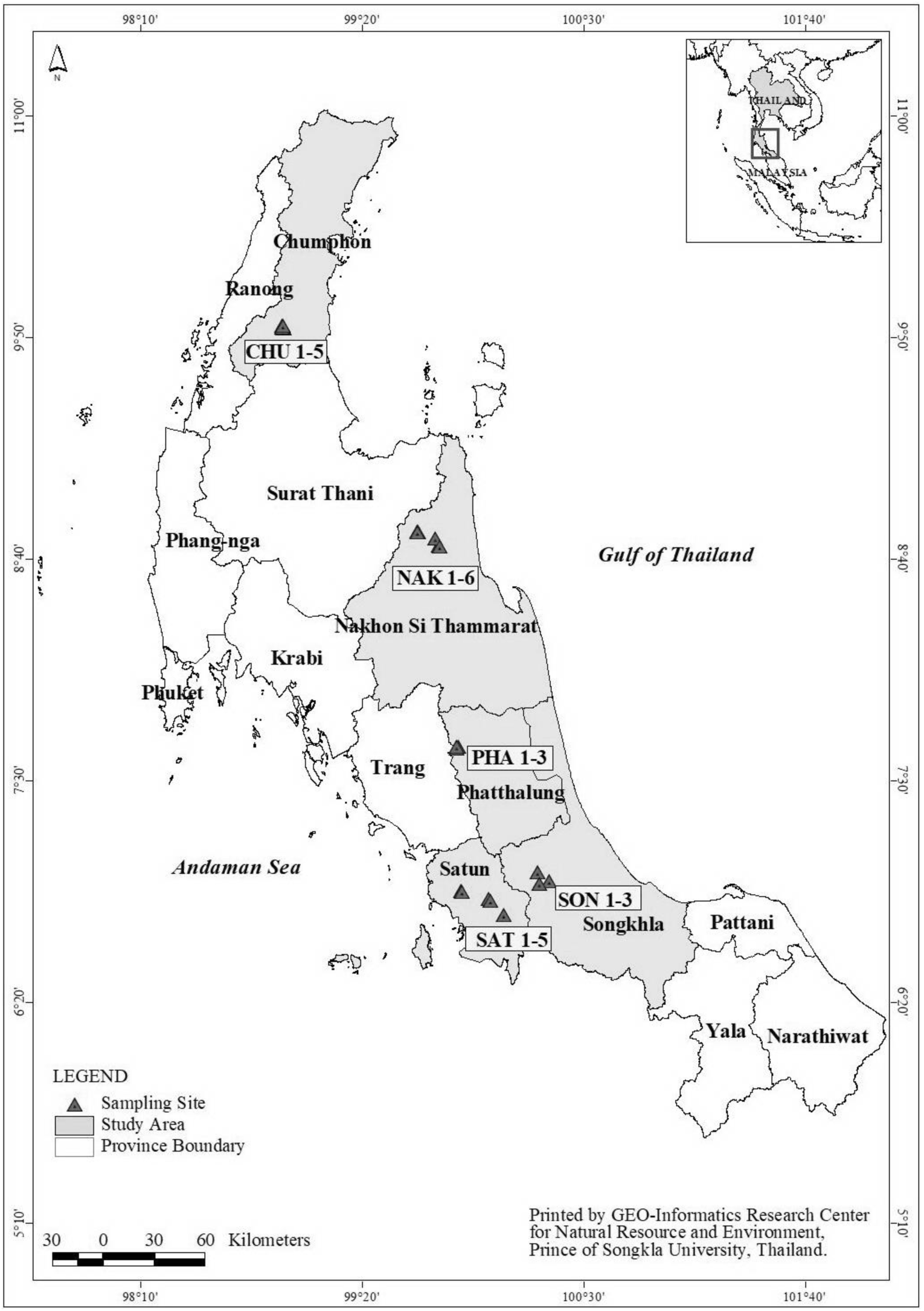
Location of durian orchards included in this study in Chumphon, Nakhon Si Thammarat, Phattalung, Satun and Songkhla provinces.

### Mosquito collection

At each study site, eggs of *Ae. albopictus*, as well as all immature stages present, were collected using ovitraps consisting of a black plastic cup of 15 cm diameter and 10 cm height lined with a piece of cotton fabric (6 x 45 cm) to provide an ovipositional site. The cup was filled with approximately 150 ml of filtered tap water and four small holes were drilled into the top of the cup to allow it to drain, especially during rainy season egg collection. At each durian orchard study site, twenty ovitraps were randomly placed on the ground, 3 m apart for a period of three to five days. Each trap was labeled with a trap number and trap position. Environmental information was observed and noted. After three to five days, the traps and the water in each trap were collected and brought back to the laboratory. The eggs on the fabric were counted and the number per trap recorded before hatching. Then, the larvae were raised at a density of 150/1,000 ml of well water in plastic trays (30 x 20 x 12 cm). The larvae were fed with fish food (Sakura, U Lek Trading Co., Ltd., Bangkok, Thailand) once a day until the pupal stage. The pupae were counted and collected daily and placed into a mesh cage to allow the adult eclosion. The adults were reared in a mesh cage (30 x 30 x 30 cm) at the Agricultural Innovation and Management Division, Prince of Songkla University under the following laboratory conditions: 25 ± 2 °C, 80 % RH and they were sustained on cotton soaked in 10 % sugar solution (in water). They were morphologically identified to species using a stereomicroscope.

### Mosquito populations used for agrochemical insecticide susceptibility test

#### *Aedes albopictus* susceptible strain

The eggs of a laboratory strain of *Ae. albopictus* were obtained from the Department of Entomology, Kasetsart University, Bangkok. This strain was originally from the Ministry of Public Health Thailand. This population had been continuously reared under laboratory conditions for over 50 generations with the adults being sustained on blood using artificial membrane feeding (Yaya and Tainchum 2017) to generate sufficient numbers of mosquitoes for insecticide susceptibility bioassays.

#### *Aedes albopictus* field populations

Immature mosquitoes collected from the orchards were mass reared as described above. Female mosquitoes aged three to five days were starved for 24 hours before insecticide susceptibility testing. Only first to fifth (F_1_-F_5_) generation females were used and mixed in tests to be representative of the field population.

#### *Aedes aegypti* susceptible strain

The eggs of a laboratory strain of *Ae. aegypti* (USDA), which originated from the Center for Medical, Agricultural, and Veterinary Entomology, Gainesville, FL, was obtained from the Department of Entomology, Kasetsart University, Bangkok. This population had been continuously reared in a laboratory for over 50 generations.

### Preparation of agrochemical insecticides

Based on the information obtained from the questionnaires regarding the type of agrochemical insecticides used in the selected durian orchards, the most frequently used agrochemical insecticides recorded were pyrethroid, organophosphate, carbamate, and neonicotinoid. The commercial form of these insecticides used against durian insect pests, along with their field application rates according to the product label, was used for bioassays. They comprised; chlorpyrifos (touchban®, 40 % EC, produced by Pro Enterprise Co., Ltd., Nakhon Chai Si, Nakhon Pathom, 60 ml/water 20 L), lambda-cyhalothrin (Karate® 2.5 EC, 2.5 % EC, produced by Syngenta Crop Protection Co., Ltd., Mueang, Samut Prakan, 25 ml/water 20 L), carbaryl (Sethrin 85®, 85 % WP, produced by Muang Thong Agriculture Co., Ltd., Lam Luk Ka, Pathumthani, 20 g/water 20 L), and imidacloprid (Pidofin®, 10 % SL, produced by SPKG Biokem Co., Ltd., Phutthamonthon, Nakhon Pathom, 10 ml/water 20 L). Tap water was used as a diluent and as a negative control.

### Insecticide-treated filter paper

Insecticide-treated papers were made at the Pest Management Laboratory, Agricultural Innovation and Management Division, Prince of Songkla University, based on the standard procedure and specifications of the World Health Organization (WHO, 2016). Insecticide-treated papers for each insecticide were prepared using Whatman® No.1, 12 x 15 cm size. The papers were treated at a rate of 2 ml of insecticide solution per sheet. Control papers were prepared in the same manner but impregnated with only 2 ml of tap water.

### WHO susceptibility tests

The insecticide susceptibility status of the *Ae. albopictus* laboratory and field strains were tested using WHO susceptibility test kits according to the WHO protocol (WHO 2016). Each set of a test kit for both treatment and control contained a pair of exposure tubes, one marked with a red dot for the insecticide-treated paper (acetone-treated paper for control) and a holding tube marked with a green dot for the untreated paper. Twenty-five three-to five-day-old, starved female *Ae. albopictus* were introduced into each respective holding tube and held for five minutes to allow the mosquitoes to adjust to the holding tubes. All the mosquitoes were subsequently exposed for 60 minutes to either treated or control paper surfaces in the exposure tubes. The number of mosquitoes knocked down in each test was recorded at 60 minutes, and all the specimens were subsequently transferred into clean holding tubes and provided with 10 % sucrose cotton pads. Four replications for each insecticide and control were performed. The mortality of the treatment and control mosquitoes were recorded after 24 hours post-exposure.

### Comparison between the susceptibility of *Aedes* mosquitoes to pyrethroid agrochemical and public health insecticides

Pyrethroid insecticide is the most commonly used public health insecticide for mosquito control and management. Two concentrations of lambda-cyhalothrin based on the agricultural application rate (0.001 g a.i. /m^2^) and the public health rate (0.01 g a.i./m^2^) were used to determine the susceptibility status of *Aedes* mosquitoes. The impregnation of the filter papers and insecticide susceptibility tests were performed as described above.

### Data analysis

Data from the questionnaires were recorded on a spreadsheet and analyzed using Microsoft Excel software (Excel® 2013). Descriptive statistics comprising means, percentages, and ranges were computed. In each study location, the patterns associated with the participant’s responses were identified. Mosquito density comparisons between each durian insecticide application system were performed using Scheffe’s multiple range test with the significance level set at *P* < 0.05. The susceptibility of mosquitoes to each insecticide was assessed. The mortality rates observed in the test and control groups were calculated according to WHO guidelines (WHO 2016).

## RESULTS

### Insecticide types and quantity used in durian planting systems in southern Thailand

As shown in Table 2, the majority (63.64 %) of the 22 durian cultivators surveyed was between 51 and 75 years old, and most (81.82 %) were male. Their highest education levels were primary, 45.45 %, secondary, 22.73 %, and Bachelor’s degree, 31.82 %, and most (90.91 %) of the respondents were farmers with the remaining 9.09 %, being government employees or officers. The form of agriculture practiced was largely polyculture (77.27 %) with 22.73 % practicing monoculture. Both forms of culture employed cultivation areas of at least 2 rai. Within the 22 orchards surveyed, the most common distance between trees was 6-10 m (81.82 %). Of all the cultivators surveyed, 68.18 % used insecticides, and the highest frequency of insecticide use per month was every 6-10 days (60.00 %), followed by 10-15 days (20.00 %), and over 15 days (20.00 %). Only 3 (13.64 %) of the durian cultivators comprising the owners of CHU 5 (IA area), SON 1 (LA area), and SON 3 (LA area), reported having been sick due to a mosquito-borne disease (dengue, chikungunya, and zika viruses) in each case having contracted dengue fever.

**Table 2.** Demographic information of durian cultivators who participated in the study.

| **Participant characteristic N (%)** |  |  |  |  |
| --- | --- | --- | --- | --- |
| **Age (years)** | **< 25** | **26-50** | **51-75** |  |
|  | 0 (0.00) | 8 (36.36) | 14 (63.64) |  |
| **Sex** | **Male** | **Female** |  |  |
|  | 18 (81.82) | 4 (18.18) |  |  |
| **Education** | **Primary school** | **Secondary school** | **Bachelor's degree** |  |
|  | 10 (45.45) | 5 (22.73) | 7 (31.82) |  |
| **Occupation** | **Farmer** | **Government employee** |  |  |
|  | 20 (90.91) | 2 (9.09) |  |  |
| **Type of durian orchard** | **Monoculture** | **Polyculture** |  |  |
|  | 5 (22.73) | 17 (77.27) |  |  |
| **Size of durian orchard (rai)** | **< 4** | **5-8** | **9-12** | **< 13** |
|  | 6 (27.27) | 7 (31.82) | 4 (18.18) | 5 (22.73) |
| **Spacing between trees (meters)** | **Indeterminate** | **0-5** | **6-10** |  |
|  | 2 (9.09) | 2 (9.09) | 18 (81.82) |  |
| **Insecticide** | **Use** | **Not use** |  |  |
|  | 15 (68.18) | 7 (31.82) |  |  |
| **Frequency of insecticide application (days)** | **1-5** | **6-10** | **11-15** | **15+** |
|  | 0 (0.00) | 9 (60.00) | 3 (20.00) | 3 (20.00) |
| **Ever had *Aedes*-borne diseases** | **Yes** | **Never** |  |  |
|  | 3 (13.64) | 19 (86.36) |  |  |

### Insecticides used to insect pest control in durian plantations

As shown in Table 3, the use of various groups of insecticides was recorded in the different durian insecticide application systems. Out of a total of 17 recorded users, a combination of organophosphate and pyrethroid insecticides was most common, accounting for 29.41 %, followed by pyrethroids (17.64 %), carbamates (17.64 %), organophosphates (11.76 %), neonicotinoids (11.76 %), pyridazinone (5.88 %) and avermectin (5.88 %). The frequency of spraying for each of these insecticides was 7-15 days per month.

**Table 3.** Type of agrochemical insecticides, their frequency of use, and the proportion of respondents who use them within the insecticide application systems included in this study.

| Insecticide group | IRAC | Active ingredient (a.i.) | Frequency of spraying (days) | Number of respondents (%) |
| --- | --- | --- | --- | --- |
| pyrethroid | A3 | cypermethrin | 15 | 1(5.88) |
|  | A3 | lambda-cyhalothrin | 15 | 2 (11.76) |
| organophosphate | B1 | chlorpyrifos | 7 | 2 (11.76) |
| organophosphate+ pyrethroid | B1+A3 | chlorpyrifos+ cypermethrin | 7, 10 | 4 (23.53) |
|  | B1+A3 | profenofos + cypermethrin | 10 | 1 (5.88) |
| carbamate | A1 | fenobucarb | 10 | 1 (5.88) |
|  | A1 | methomyl | 7 | 1 (5.88) |
|  | A1 | carbaryl | 15 | 1 (5.88) |
| neonicotinoid | A4 | imidacloprid | 10, 15 | 2 (11.76) |
| pyridazinone* |  | pyridaben | 7 | 1 (5.88) |
| avermectin | 6 | abamectin | 10 | 1 (5.88) |

Insecticide grouping is based on the Insecticide Resistance Action Committee (IRAC) classification
\*Herbicide

### Determining the density of *Ae. albopictus* in durian planting systems of southern Thailand

Figure 2 shows the number of mosquito eggs per trap along with the Scheffe multiple range test results comparing the number of eggs per trap between orchards. In the three durian insecticide application systems, IA, LA, and NA, the mean number of eggs per trap ranged from 4.40-63.70, 10.00-50.35 and 6.16-115.20, respectively, and significant differences between durian plantations (*P* < 0.05 were found as shown in Figure 2. The site with the most mosquito eggs was PHA 1 (115.20 ± 12.83) followed by PHA 2 (73.25 ± 21.49) among the NA classification, and PHA 3 (63.70 ± 10.69) among the IA classification. However, no mosquito eggs were collected from 58.33 % of the IA orchards. In addition, the mean number of pupae collected from the three durian insecticide application systems, IA, LA, and NA, were in the range of 2.05-26.20, 1.42-39.80, and 10.05-39.60, respectively. The three durian cultivation sites with the highest number of pupae were SAT 4 (26.20), SON 1 (39.80), and SAT 5 (39.60). The three sites with the lowest numbers of pupae were PHA 3, SON 3, and SON 2. All of the eggs collected from NAK 5 and PHA 2 either failed to hatch or did not develop to the pupal stage (Figure 3).

**Figure 2.**
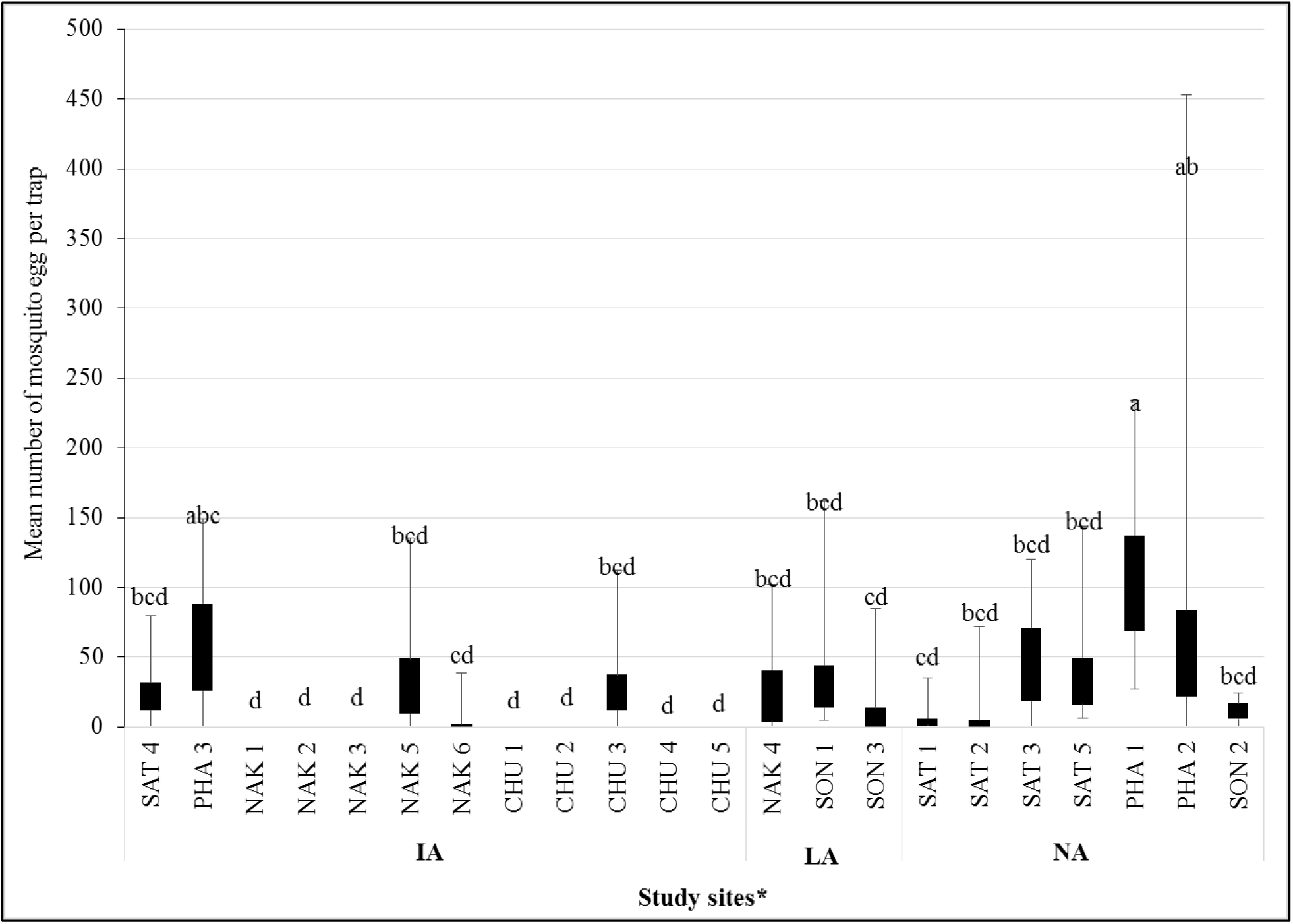
Mean number of *Aedes albopictus* eggs/ovitraps in each study site and Scheffe’s multiple range test between each orchard *IA = intensive-application of insecticides, LA = less-application of insecticides, and NA = no application of insecticides, the same letters (a-d) are non-significantly different at *P*>0.05.

**Figure 3.**
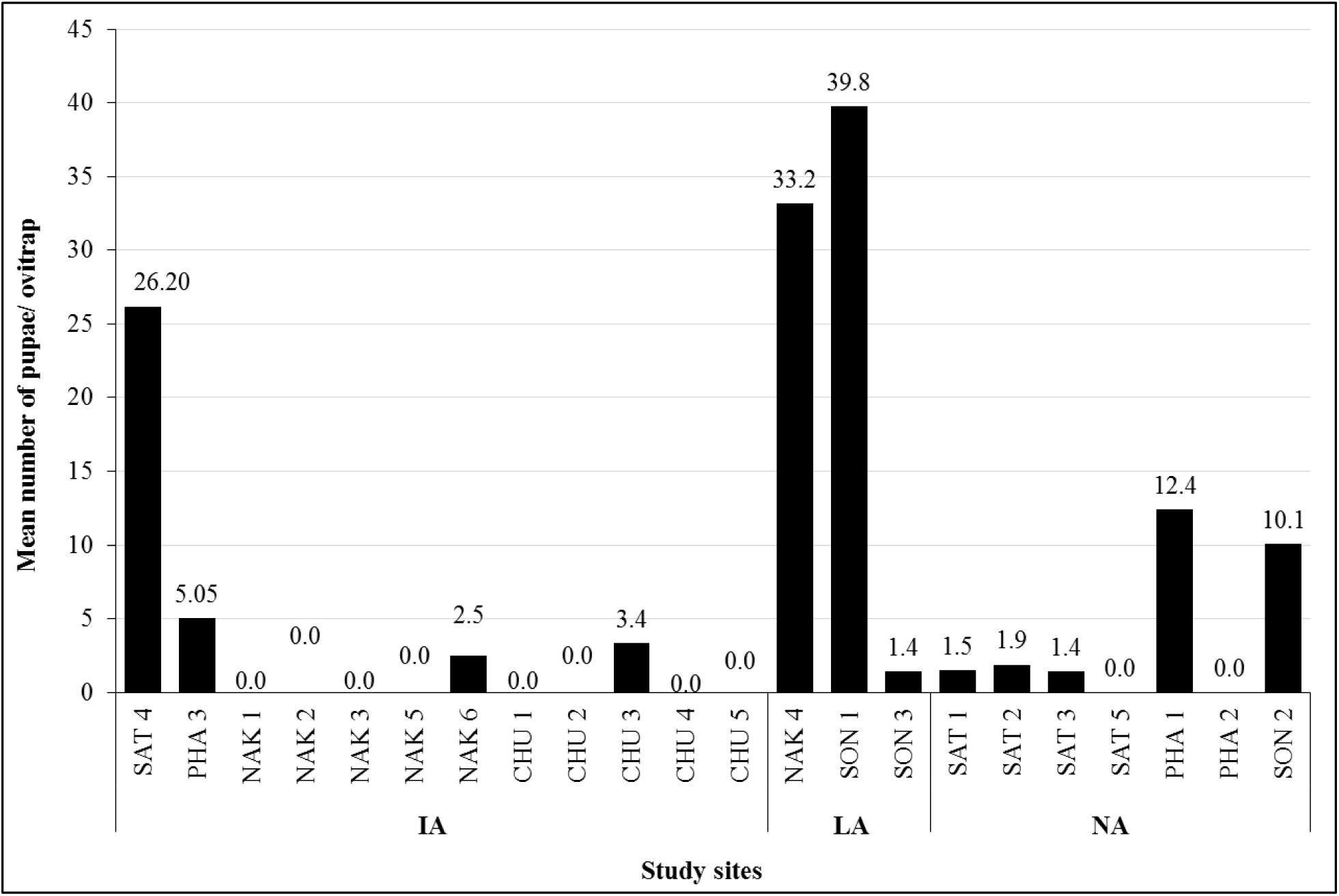
Mean number of *Aedes albopictus* pupae per study site.

### Susceptibility of *Ae. albopictus* to frequently used agrochemical-insecticides in durian planting systems

The susceptibility tests conducted on laboratory strains of *Ae. aegypti* and *Ae. albopictus*, as well as on field-collected *Ae. Albopictus* mosquitoes, revealed variation in the proportions of knockdown insects and mortality between both insecticides and study sites. The proportion of laboratory (NIH) and field strains of *Ae. albopictus* knocked down, as well as laboratory *Ae. aegypti* strain (USDA) following 60 minutes of exposure to field application concentrations of chlorpyrifos, lambda-cyhalothrin, carbaryl, and imidacloprid are shown in Table 4. The mosquitoes used as controls were all alive after bioassay with 0 % knockdown. Overall the percentage knockdown from highest to lowest percentage was imidacloprid < carbaryl < lambda-cyhalothrin < chlorpyrifos. Surprisingly, a high proportion of those knockdown was observed in both species of laboratory strains (5-100 % knockdown), for all the insecticides except imidacloprid. Less than 3 % of those knockdowns were recorded in the field population of *Ae. albopictus* exposed to imidacloprid at SAT 4 (1.25 %) in an IA site, and PHA 1 (2.50 %) in an NA site. The remaining populations were completely knockdown. All of the mosquito populations that were exposed to chlorpyrifos were 100 % knockdown, except for PHA 3 (76 %) which was an IA site. The percentage mortality of the laboratory (NIH) and field strains of *Ae. albopictus*, as well as of the laboratory strain of *Ae. aegypti* (USDA) after 24 hours of exposure to field application concentrations of chlorpyrifos, lambda-cyhalothrin, carbaryl, and imidacloprid is shown in Table 5. There was no mortality in any of the controls. Complete (100 %) mortality was seen in all populations after exposure to chlorpyrifos from the organophosphate insecticide group. For the pyrethroids, the proportion of mortality recorded in all the populations following 24 hours exposure to lambda-cyhalothrin ranged between 46.23 and 81.20 %, except in PHA 3, which was an IA site, and showed higher mortality (96.84 %). Among the carbamates, most of the populations exposed to carbaryl recorded mortality below 90 % (mortality range, 40.00-88.73 %) although the highest mortality was recorded at SON 1 (96.05 %), which was an LA site. In the neonicotinoid group, all the mosquito populations were exposed to imidacloprid, and all the mortality rates recorded were below 11 % (0.00-10.33 %; see Table 5).

**Table 4.** Percentage mosquitoes knockdown among laboratory and field strains of *Aedes albopictus*, and laboratory strain of *Aedes aegypti* following 24 h exposure to field application concentrations of chlorpyrifos, lambda-cyhalothrin, carbaryl and imidacloprid.

| Population | Control | Chlorpyrifos | Lambda-cyhalothrin | Carbaryl | Imidacloprid |
| --- | --- | --- | --- | --- | --- |
| USDA<sup>*</sup> | 0.00 | 100.00 | 80.00 ± 4.33 | 26.67 ± 17.02 | 0.00 |
| NIH | 0.00 | 100.00 | 76.25 ± 17.37 | 5.00 ± 4.33 | 0.00 |
| **IA** |  |  |  |  |  |
| SAT 4 | 0.00 | 100.00 | 41.25 ± 8.51 | 2.50 ± 2.50 | 1.25±1.25 |
| PHA 3 | 0.00 | 76.00 ± 4.62 | 91.00 ± 4.12 | 2.00 ± 1.15 | 0.00 |
| **LA** |  |  |  |  |  |
| NAK 4 | 0.00 | 100.00 | 68.75 ± 3.75 | 18.33 ± 15.88 | 0.00 |
| SON 1 | 0.00 | 100.00 | 65.65 ± 4.72 | 53.47 ± 12.91 | 0.00 |
| **NA** |  |  |  |  |  |
| SAT 5 | 0.00 | 100.00 | 80.00 ± 7.36 | 7.50 ± 2.95 | 0.00 |
| PHA 1 | 0.00 | 100.00 | 47.50 ± 4.79 | 7.50± 3.23 | 2.50 ± 1.44 |

IA = intensive-application of insecticides, LA = less-application of insecticides, and NA = no application of insecticides
<sup>\*</sup>Laboratory strain, USDA = *Ae. aegypti* and NIH = *Ae. albopictus*

**Table 5.** Percentage mortality of laboratory and field strains of *Aedes albopictus*, and laboratory strain of *Aedes aegypti* following 24 h exposure to field application concentrations of chlorpyrifos, lambda-cyhalothrin, carbaryl and imidacloprid.

| Populations | Control | Chlorpyrifos | Lambda-cyhalothrin | Carbaryl | Imidacloprid |
| --- | --- | --- | --- | --- | --- |
| USDA* | 0.00 | 100.00 | 62.11 ± 15.70 | 40.00 ± 24.62 | 0.00 |
| NIH | 0.00 | 100.00 | 65.03 ± 8.53 | 57.21 ± 15.83 | 0.00 |
| IA |  |  |  |  |  |
| SAT 4 | 0.00 | 100.00 | 77.35 ± 4.10 | 56.62 ± 13.12 | 1.25 ± 1.25 |
| PHA 3 | 0.00 | 100.00 | 96.84 ± 2.16 | 61.29 ± 5.21 | 0.00 |
| LA |  |  |  |  |  |
| NAK 4 | 0.00 | 100.00 | 81.20 ± 3.45 | 52.11 ± 12.24 | 0.00 |
| SON 1 | 0.00 | 100.00 | 46.23 ± 10.68 | 96.05 ± 3.95 | 0.00 |
| NA |  |  |  |  |  |
| SAT 5 | 0.00 | 100.00 | 77.31 ± 5.89 | 80.15 ± 12.89 | 1.25 ± 1.25 |
| PHA 1 | 0.00 | 100.00 | 59.38 ± 8.64 | 88.73 ± 7.86 | 10.33 ± 4.56 |

IA = intensive-application of insecticides, LA = less-application of insecticides, and NA = no application of insecticides
\*Laboratory strain, USDA = *Ae. aegypti* and NIH = *Ae. albopictus*

### Comparison between the susceptibility of *Aedes* mosquitoes to pyrethroid agrochemical and public health insecticides

An initial study comparing the susceptibility of *Aedes* mosquitoes between application concentrations of lambda-cyhalothrin for agrochemical (AL) and public health (PL) use, showed a higher overall proportion of mosquitoes knockdown (> 94.80 %) and mortalities (96.15 %) in all the *Aedes* populations that were exposed to lambda-cyhalothrin as applied as a public health insecticide compared to its application as an agrochemical (knockdown = 37.84-97.50 % and mortalities = 45.75-86.43 %). For the field strain of *Ae. albopictus*, a higher proportion of mosquitoes knockdown and mortality were seen at its public-health application dosage compared to its dosage as an agrochemical (Table 6).

**Table 6.** Comparison between the susceptibility of *Aedes* mosquitoes to agrochemical and public health application dosages of lambda-cyhalothrin.

| Population | %KD 60 minutes |  |  | %Mortality 24 hours |  |  |
| --- | --- | --- | --- | --- | --- | --- |
|  | Control | PL** | AL*** | Control | PL | AL |
| USDA* | 0.00 | 100.00 | 66.67 | 0.00 | 100.00 | 50.00 |
| NIH | 0.00 | 100.00 | 37.84 | 0.00 | 100.00 | 45.75 |
| IA |  |  |  |  |  |  |
| SAT 4 | 0.00 | 96.15 | 92.09 | 10.00 | 98.72 | 82.34 |
| PHA 3 | 0.00 | 100.00 | 88.00 | 0.00 | 100.00 | 76.00 |
| LA |  |  |  |  |  |  |
| NAK 4 | 0.00 | 94.87 | 82.59 | 0.00 | 100.00 | 78.95 |
| SON 2 | 0.00 | 100.00 | 92.17 | 47.50 | 98.68 | 70.62 |
| NA |  |  |  |  |  |  |
| SAT 5 | 2.50 | 100.00 | 97.50 | 10.00 | 97.30 | 86.43 |
| PHA 1 | 0.00 | 98.72 | 84.87 | 5.00 | 96.15 | 56.98 |

IA = intensive-application of insecticides, LA = less-application of insecticides, and NA = no application of insecticides
\*Laboratory strain, USDA = *Ae. aegypti* and NIH = *Ae. albopictus*
\*\*PL = lambda-cyhalothrin used as a public health insecticide for mosquito control.
\*\*\*AL = lambda-cyhalothrin used as an agricultural insecticide for the control of target insect pests.

## DISCUSSION

The objective of this study was to observe the density of *Ae. albopictus* in durian planting systems in southern Thailand and to evaluate their insecticide-resistance status. The small durian farmers who took part in the study were, however, not necessarily representative of durian cultivators in southern Thailand. A similarly designed study was conducted by de Albuquerque et al. (2018) in which ovitraps were set for 15 or 30 days near a house in the urban areas of Itacoatiara and Tabatinga, in Amazonas, Brazil to examine the density of *Ae. aegypti.* That study found a positive correlation between the occurrence of dengue and the *Ae. aegypti* egg density. Previous work by Regis et al. (2008) on the density of *Ae. albopictus* determined using ovitraps placed near forested areas with high rates of disease transmission, showed that the *Aedes* egg density index (EDI) is equal to 100-750 eggs per trap.

Ovitraps for mosquito collection follow many designs and can be made from various kind of material. The ovitraps used in this study followed the well-known ovitrap design launched by the Center for Disease Control and Prevention in the USA, which can be made using a small metal, glass, or plastic container, often dark in color, containing water and material in which females can lay eggs. This trap, which is inexpensive and easily transportable, is mainly used to survey the population of *Aedes* mosquitoes. One drawback of the use of ovitraps is that they may become mosquito breeding sites if left for more than a week. Additionally, environmental and/or human activities may contribute mosquito breeding sites that may compete with ovitraps, thus compromising the number of eggs collected by an ovitrap (CDC 2018).

The insecticides used in this experiment, organophosphate (chlorpyrifos), pyrethroid (lambda-cyhalothrin), carbamate (carbaryl) and neonicotinoid (imidacloprid) were applied based on the recommended concentrations on the labels and produced active ingredient per square meter (a.i./m^2^) levels of 0.04, 0.001, 0.03 and 0.002 g, respectively. However, the concentrations of insecticides recommended by the WHO for public health use for mosquito control are as follows; organophosphate (fenitrothion) 2.0 g a.i./m^2^, pyrethroid (lambda-cyhalothrin) 0.02-0.03 g a.i./m^2^ and carbamate (propoxur) 1.0-2.0 g a.i./m^2^, with neonicotinoids not having yet been approved for public health use (WHO 2015). Therefore the a.i./m^2^ recommended for agricultural purposes is much less than that approved for use in public health applications. Since mosquitoes are non-target insects for agricultural insecticides, continued exposure to sub-lethal concentrations of agricultural insecticides could select for insecticide resistance in mosquito populations. This is the probable cause of the reduced mortality in the mosquito populations tested in this study for all the insecticides except chlorpyrifos.

However, the low knockdown and mortality rates in the mosquito populations tested may not be due to their lower insecticide susceptibility. For example imidacloprid is an insecticide in the neonicotinoid group, all of which are synthetic substances which imitate the action of nicotine. The mode of action of this insecticide group is to bind to the nicotinic acetylcholine receptor in the central nervous system, thus blocking signal transmission to nerve cells. Imidacloprid enters the insect’s system by being eaten (Gervais 2010) but in this bioassay, which employed the WHO susceptibility test based on tarsal contact, the neonicotinoid was not able to enter the mosquitoes’ system in order to act. Therefore it cannot be concluded that the mosquitoes in this study were resistant to imidacloprid.

The results of the comparisons between the susceptibility of *Aedes* mosquitoes to agrochemical and public health lambda-cyhalothrin insecticides showed that *Ae. albopictus* from the study sites were mostly susceptible to the public health dosage of lambda-cyhalothrin, with overall mortality of 96.15 %. This was in contrast to a mortality of 86.43 % for the agrochemical dosage of lambda-cyhalothrin. Further, our results showed no evidence of cross-insecticide resistance between agrochemical and public health lambda-cyhalothrin insecticides. In the future, cross-resistance between agrochemical and public health insecticides (organophosphate, pyrethroid, or carbamate) should be a required component of insecticide resistance management.

Overall, the populations of *Ae. albopictus* in this study were completely susceptible to chlorpyrifos but experienced reduced mortality following exposure to lambda-cyhalothrin, carbaryl, and imidacloprid, which is suggestive of the existence of resistance. To the best of our knowledge, this is the first report of susceptibility tests in respect of agrochemical insecticides on wild populations of *Aedes* in southern Thailand. Previous studies have however reported the insecticide susceptibility status of *Aedes* mosquitoes against recommended public health concentrations of insecticides for vector control. Thanispong et al. (2008) reported that *Ae. aegypti* from Muang district, Songkhla province and Muang district, Satun province exposed to the recommended public health concentration of alpha-cypermethrin (0.05 %), deltamethrin (0.05 %), permethrin (0.25 %), and malathion (0.8 %) were both susceptible to deltamethrin, malathion, and alpha-cypermethrin. However, *Ae. aegypti* in Songkhla showed sime suggestion of resistance to alpha-cypermethrin and also to permethrin. In a later study by Chuaycharoensuk et al. (2011), *Ae. albopictus* in rubber plantations from Songkhla and Chumphon provinces were susceptible to deltamethrin and lambda-cyhalothrin, while the Chumphon strain exhibited some suggestion of resistance to permethrin and the Songkhla stain were resistant to permethrin.

## CONCLUSION

The most commonly used groups of insecticides in durian plantations in the five provinces in southern Thailand (Chumphon, Nakhon Si Thammarat, Phattalung, Satun and Songkhla) were: organophosphate combined with a pyrethroid (chlorpyrifos + cypermethrin), followed by pyrethroid (cypermethrin and lambda-cyhalothrin), carbamate (fenobucarb, methomyl, and carbamate), organophosphate (chlorpyrifos) and neonicotinoid (imidacloprid). Frequent applications (7-15 days per month) of each insecticide for insect pest control were recorded for more than half the sample. The variation in insecticide intensity and frequency in durian plantations influenced the density of *Ae. albopictus* eggs collected by ovitraps, as well as disrupting the mosquito life cycle by hindering the adult female mosquitoes from completing their gonotrophic cycle, and thus egg-laying. The number of eggs collected was significantly different (*P* < 0.05) among the three durian plantation insecticide application criteria IA, LA and NA. Unsurprisingly, the highest number of eggs per trap was collected from the NA sites in which no insecticides were used, followed by the LA and IA sites, respectively.

Of the four groups of insecticides used in the durian plantations in this study three are also used in public health applications for vector control: organophosphate (chlorpyrifos), pyrethroids (lambda-cyhalothrin, cypermethrin) and carbamate (carbaryl), but at different concentrations, resulting in different dosages of active ingredients. Their use in durian farming may lead to the development of insecticide resistance in mosquito populations, as well as cross-resistance to public health insecticides. However, since the mosquitoes in this study were completely susceptible to chlorpyrifos, should other insecticides fail, that appears to be a good alternative for *Ae. albopictus* control.

Finally, the monitoring of insecticide susceptibility and the early detection of insecticide resistance should always be considered in the design and implementation of effective integrated vector management practices for the control of *Aedes*-borne diseases in Thailand.

## Acknowledgments

This research study was funded by a Research Grant for Thesis, from the Graduate School, and a Graduate Thesis Grant from the Faculty of Natural Resources, Prince of Songkla University. The authors are grateful to the durian orchard cultivators, without whose cooperation this study would not have been possible.

